# Comparing Methods for Species Tree Estimation With Gene Duplication and Loss

**DOI:** 10.1101/2021.02.05.429947

**Authors:** James Willson, Mrinmoy Saha Roddur, Tandy Warnow

## Abstract

Species tree inference from gene trees is an important part of biological research. One confounding factor in estimating species trees is gene duplication and loss which can lead to gene trees with multiple copies of the same gene. In recent years there have been several new methods developed to address this problem that have substantially improved on earlier methods; however, the best performing methods (ASTRAL-Pro, ASTRID-multi, and FastMulRFS) have not yet been directly compared. In this study, we compare ASTRAL-Pro, ASTRID-multi, and FastMulRFS under a wide variety of conditions. Our study shows that while all three have very good accuracy, nearly the same under many conditions, ASTRAL-Pro and ASTRID-multi are more reliably accurate than FastMuLRFS, and that ASTRID-multi is often faster than ASTRAL-Pro. The datasets generated for this study are freely available in the Illinois Data Bank at https://databank.illinois.edu/datasets/IDB-2418574

## 1 Introduction

Species tree estimation is an important part of biological research. One of the main challenges is gene tree heterogeneity, when the individual genes of a species evolve differently and thus the topology of the gene family trees does not match that of the species tree. This can be caused by several phenomena, such as gene duplication and loss (GDL), incomplete lineage sorting (ILS), horizontal gene transfer, and hybrid speciation. There are many methods available to estimate species trees in the presence of ILS that are proven statistically consistent under the Multi-Species Coalescent [6] model, and many of these have very good accuracy on large datasets (e.g., ASTRAL [10], ASTRID [18]). However GDL is another of the primary causes of gene tree heterogeneity, and so methods that can estimate species trees with high accuracy in the presence of GDL are needed.

One approach to species tree estimation in the presence of GDL presumes the ability to identify orthologs (i.e., genes in different species that have evolved from a common ancestor via a speciation rather than a duplication event). Given the accurate identification of orthologs, species tree estimation can proceed by restricting analyses to single-copy gene trees. However, orthology detection is not yet reliably solved [5], and so this approach can result in errors in the input (i.e., the single copy gene trees may not reflect the species tree), and hence errors in the output species tree. Furthermore, restricting to orthologous genes reduces the amount of available phylogenetic signal, and so can reduce accuracy.

An alternative approach is to use the multi-copy genes (either their alignments or their gene family trees, called “MUL-trees”), and then use these to estimate the species tree. Methods that take this approach include PHYLDOG [2] and guenomu [13], which use probabilistic models of gene evolution, and so are comparatively slow. Gene tree parsimony approaches (seeking to minimize the total number of duplications and losses, NP-hard problems) have also been developed, including DupTree [19], iGTP [3], and DynaDup [1]. None of these methods have yet been proven to be statistically consistent under GDL models.

In 2019, a species tree estimation method that was developed for the ILS-only scenario, ASTRAL-multi [15], was proven to be statistically consistent under GDL [8]. Subsequently, ASTRAL-Pro [20] was developed to explicitly address GDL; its technique requires “rooting and tagging” each input gene tree, and if the rooting and tagging is correctly performed, then ASTRAL-Pro is statistically consistent under GDL. Interestingly, although ASTRAL-Pro has not been proven statistically consistent under GDL, it has been shown to be more accurate than ASTRAL-multi in extensive simulation studies [20].

ASTRID-multi (a recent improvement of the ASTRID [18] method for species tree estimation under ILS) and FastMulRFS [11] (a method for the Robinson-Foulds Supertree problem adapted to the MUL-tree setting) have also been found to be more accurate than ASTRAL-multi in simulation studies evaluating methods for species tree estimation under models that include GDL and varying levels of ILS [8, 11, 20]. However, a comparison between these three leading methods has not been performed, and no analyses have been made of these methods when gene trees are missing species (i.e., the “missing data” condition), which is very common in biological datasets.

Here, we report on a study comparing ASTRAL-Pro, ASTRID-multi, and FastMulRFS in terms of topological accuracy and running time on data sets we simulated under the DLCOAL model [16] that includes GDL and ILS. Unlike previous studies, we include conditions where gene trees are “incomplete”, and so are missing species, and we explore performance under a wide range of model conditions, including large numbers of species (up to 1000) and genes (up to 10,000). We find that differences in accuracy between the three methods tend to be small, but that there are conditions where FastMulRFS is not quite as accurate as either ASTRAL-Pro or ASTRID-multi. The comparison between ASTRID-multi and ASTRAL-Pro shows very close accuracy under most conditions and that ASTRID-multi is faster. Overall, we find that ASTRID-multi and ASTRAL-Pro are both reliable methods for species tree estimation from multi-locus datasets that do not require orthology determination, and that of the two, ASTRID-multi is faster.

## 2 Experiment Design

To compare ASTRAL-Pro, FastMulRFS, and ASTRID-multi, we designed a simulation study with three experiments, each containing both GDL and ILS.

– Experiment 1: we examine how the methods perform on complete gene trees (i.e., no missing data) under various conditions, varying degree of GDL and ILS, relative probabilities of gene duplication and loss, sequence length per gene, and number of genes.
– Experiment 2: we examine the methods on gene family trees where some gene trees can be incomplete (i.e., the missing data condition).
– Experiment 3: we evaluate the methods scalability by running them on a simulated data set with 1000 species.

The simulation study evolved gene trees within species trees using SimPhy [9], and then evolved sequences within the gene trees using INDELIBLE [4] under the GTRGAMMA model of sequence evolution. Gene trees were then estimated on these sequence alignments using FastTree2 [14], a fast maximum likelihood heuristic, and provided as input to the different species tree estimation methods.

We ran all three methods on all the data sets and compared them with respect to species tree error and running time. For species tree error, we report the Robinson-Foulds [17] error rate between the true species tree and the estimated species tree. For Experiments 1 and 2, we ran the methods on the University of Illinois at Urbana-Champaign Campus Cluster, which imposes a four hour restriction; for Experiment 3 we use the Tallis queue which removes this restriction and has more available memory. In each experiment, we note which methods failed and for what reason (e.g., failure to complete within the allowed time, insufficient memory, or other failure).

In our simulations, we varied the number of species (from 100 to 1000), the number of genes (from 100 to 10,000), the degree of ILS, the GDL rate, the relative frequency of duplications and losses, and the gene tree estimation error. While varying a given condition, we kept the rest of the conditions locked to the following default values: 100 species, 1000 or 10,000 gene trees estimated from 100bp alignments, haploid effective population size of 5.0 *×* 10^7^, duplication rate 5.0 *×* 10^*−*10^, and a loss rate equal to the duplication rate. Thus, our default conditions had moderate levels of mean gene tree estimation error (here, 43% MGTE) and moderate levels of ILS (average topological distance between true gene trees and true species trees of 20%, denoted by AD=20%). To ensure reproducibility, all datasets simulated for this study are available upon request, and will be made freely available (after acceptance) through the Illinois Data Bank, provided through the University of Illinois. Commands necessary to reproduce the experiment are provided in the Supplementary Materials, provided at http://tandy.cs.illinois.edu/gdl-suppl.pdf.

### Experiment 1

We simulated a set of 100-taxon data sets which varied the following conditions:

#### Gene Duplication and Loss (GDL)

We chose duplication rates of 1.0 *×* 10^*−*10^, 5.0 *×* 10^*−*10^, and 1.0 *×* 10^*−*9^, all run with relative loss rates of 0, 0.5, and 1, giving us a total of nine conditions. As seen in Table 1, this gives us a set of model conditions that vary substantially in terms of average tree size (number of leaves) in the resulting gene family trees, ranging from 117 leaves to 3728 leaves.

**Table 1.** Average number of leaves in the true gene family trees for Experiment 1 (100 taxa), for different duplication rates (rows) and ratios of losses to duplications (columns). Results shown are averaged across 1000 genes and replicates.

|  | L/D=0 | L/D=0.5 | L/D=1 |
| --- | --- | --- | --- |
| $1 \times 10^{-10}$ | 145.1 | 128.0 | 116.6 |
| $5 \times 10^{-10}$ | 550.0 | 290.6 | 165.3 |
| $1 \times 10^{-9}$ | 3727.8 | 993.0 | 228.5 |

#### Number of gene trees

The number of genes is known to impact accuracy, with better accuracy obtained with more genes; therefore, we tested data sets with 100, 500, 1000, and 10,000 gene trees.

#### Mean Gene Tree Estimation Error (MGTE)

We explore results using both true and estimated gene trees. It is well known that species tree estimation methods that operate by combining gene trees are impacted by gene tree estimation error [8, 11, 20]. To explore the impact of this on the relative and absolute performance of the species tree methods we examine, we varied the sequence length provided to FastTree2 from 100 to 500 bp. This produced datasets with mean gene tree error (MGTE) of 43% and 19.2%, respectively, where gene tree error is computed using Robinson-Foulds (RF) error rates.

#### Incomplete Lineage Sorting (ILS)

We controlled the level of ILS through the effective haploid population size, choosing sizes of 1.0 *×* 10^4^, 5.0 *×* 10^7^, and 2.0 *×* 10^8^. These produce true gene trees that have average topological distance (AD) using normalized RF distance to the true species tree of 0.000468%, 20.3019%, and 50.00392%, respectively; we round these values to 0% (low ILS), 20% (moderate ILS) and 50% (high ILS) for the sake of brevity.

### Experiment 2

We generated datasets where some gene trees are missing species, using the *M*_*clade*_ model of missing data described in Nute et al. [12]. First, we listed all the clades from the species tree where the number of leaves at least 20% of all the leaves in the tree. Then, for each gene tree, we picked one of previously listed clades at random and deleted all the taxa from that gene tree which did not belong to the selected clade. On average, this protocol deleted 40% of the species from the gene trees. We ran this process on the 100-taxon data set from Experiment 1 with 100, 500, and 1000 estimated gene trees.

### Experiment 3

We simulated an additional data set with 1000 species, where the rest of the conditions were identical to the default conditions in Experiment 1. This experiment was run on the Tallis cluster, which allows the methods an unlimited amount of time to run and extra available memory.

## 3 Results

### Experiment 1

Our first analysis examined the impact of changing the gene tree estimation error on species tree accuracy (Figure 1). Given true gene trees, ASTRAL-Pro produced the true species tree while the other methods had some error (but very low error, at most 2%). On estimated gene trees, species tree error rates increased for all methods, with FastMulRFS slightly worse than the other methods, but the differences between methods were small. For running time, FastMulRFS had the highest running time, followed by ASTRAL-Pro, and then by ASTRID-multi. Furthermore, ASTRID-multi’s running time seemed completely unaffected by the change in MGTE, but FastMulRFS and ASTRAL-Pro both increased in running time as MGTE increased.

**Fig. 1.**
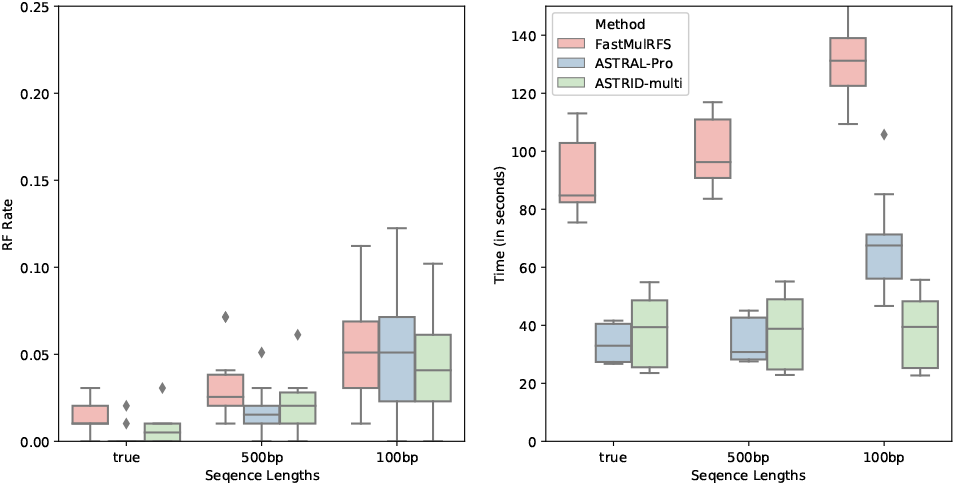
*Experiment 1*. Impact of gene tree estimation error (MGTE) on tree error (RF error rates) and wall clock running time (seconds); averages over 10 replicates per model condition are shown. All the data sets have 100 species, 1000 gene trees, AD=20%, a duplication rate of 5.0 *×* 10^*−*10^ and an equal loss rate. The data sets have MGTE of 0%, 19.2%, and 43.2%, respectively. The default conditions are the 100bp set, found furthest to the right. The boxes stretch from the 1st to 3rd quartile and the lines through the boxes shows the median.

The results on the data sets with varying levels of ILS (Figure 2) show that increasing ILS results in increased error for all methods, but the methods are comparable in accuracy until the high ILS condition (AD=50%), where FastMulRFS has a substantial increase in error rates, while the other methods remain fairly low in error. The running time shows an interesting trend: as the amount of ILS increases the running times for all methods increase, except for ASTRID-multi which remains unaffected. Also, FastMulRFS is the slowest of the methods, ASTRAL-Pro is intermediate, and ASTRID-multi is the fastest.

**Fig. 2.**
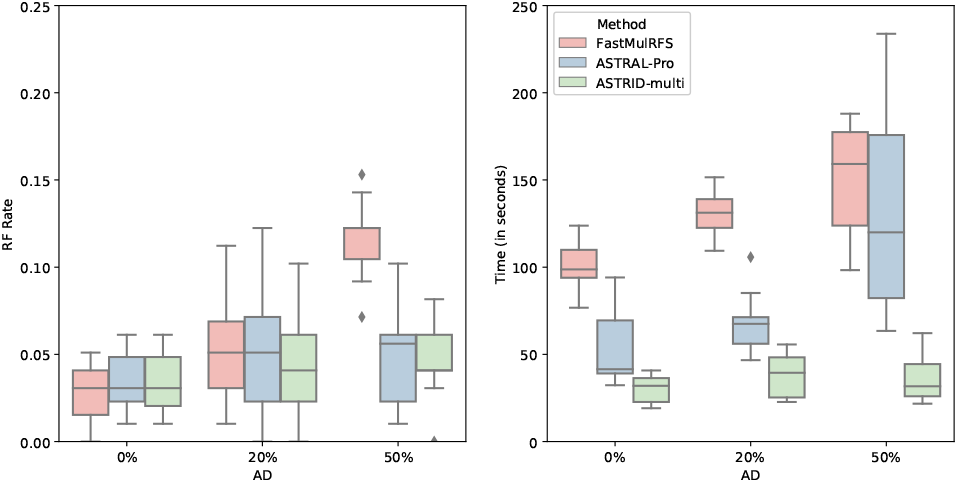
*Experiment 1*. Impact of ILS level on tree error (RF error rates) and wall clock running time (seconds); averages across 10 replicates per model condition are shown. All the data sets have 100 species, 1000 gene trees estimated from 100bp alignments (34.7%, 43.2%, 43.9% MGTE), a duplication rate of 5.0 *×* 10^*−*10^, and an equal loss rate.

We then examined the results for varying the duplication and loss rates (Figure 3). For the most part, the methods stayed competitive at lower duplication rates, but as the duplication rates increased FastMulRFS’s error rate increased, making it clearly less accurate than the others; this trend held for all duplication/loss ratios tested. The running time revealed interesting trends. For the lowest duplication rate, ASTRID-multi was the fastest and FastMulRFS the slowest, but at the highest duplication rate they reversed positions.

**Fig. 3.**
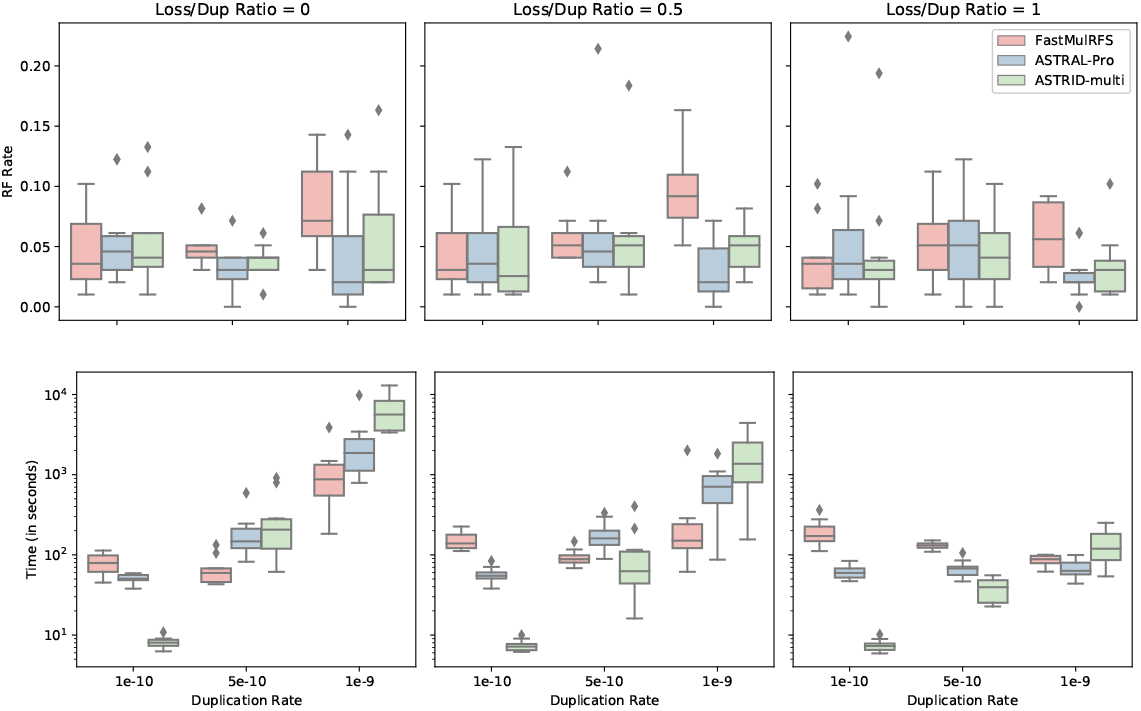
*Experiment 1*. Impact of GDL rate on tree error (RF error rates) and wall clock running time (seconds); averages across 10 replicates are shown. All the data sets have 100 species, 1000 gene trees, gene trees estimated from 100bp alignments (44.1%, 43.1%, 47.8%, 44.9%, 45.1%, 43.5%, 44.0%, 43.2%, 40.0% MGTE), and AD=20%. ASTRAL-Pro and ASTRID-multi failed on some replicates; results shown here are for the replicates on which all methods completed.

We next examined performance on datasets containing 10,000 gene trees and varying the ratio of duplication to loss (Figure 4). The three methods were indistinguishable with respect to accuracy on these data, but there were substantial differences in running time that depended on the ratio of duplications to losses. In the duplication-only scenario, FastMuLRFS was by far the fastest and the two other methods were about the same. For the other conditions, ASTRID-multi was the fastest of the three methods, and the relative running time between FastMulRFS and ASTRAL-Pro depended on the condition.

**Fig. 4.**
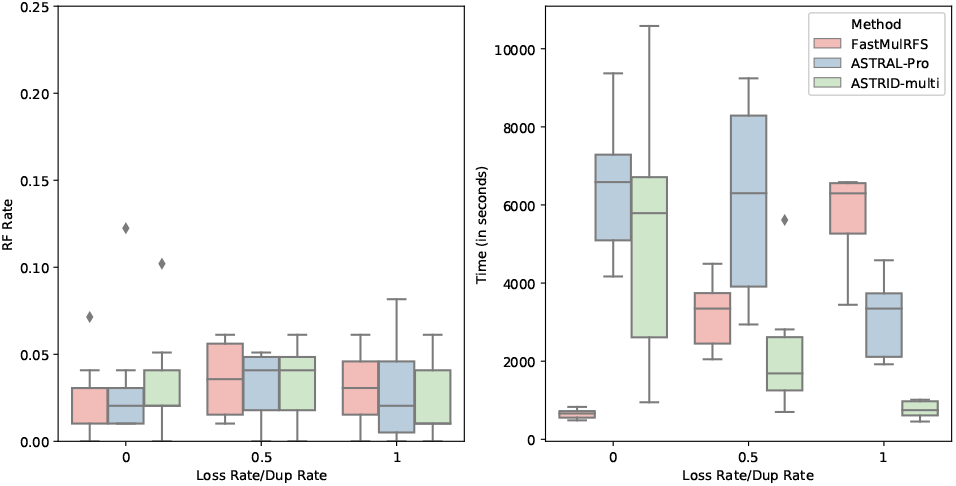
*Experiment 1*. Impact of loss rate on tree error (RF error rates) and wall clock running time (seconds) on the data sets with 10,000 gene trees; averages over 10 replicates are shown. All the data sets have 100 species, gene trees estimated from 100bp alignments (30.0%, 43.8%, 41.2% MGTE), a duplication rate of 5.0 *×* 10^*−*10^, and AD=20%. ASTRAL-Pro and ASTRID-multi failed on some replicates; results shown here are for the replicates on which all methods completed.

Next we examined the data sets which varied the number of gene trees (Figure 5). All methods increased in accuracy and running time as the number of gene trees increase. The differences between methods are largest at 100 genes, where FastMulRFS has lower accuracy than the other methods, and then reduce as the number of genes increase, so that by 1000 genes, the differences seem minor. Running time, however, is significantly impacted by the number of genes, with ASTRID-multi always faster than the other methods and FastMulRFS always the slowest.

**Fig. 5.**
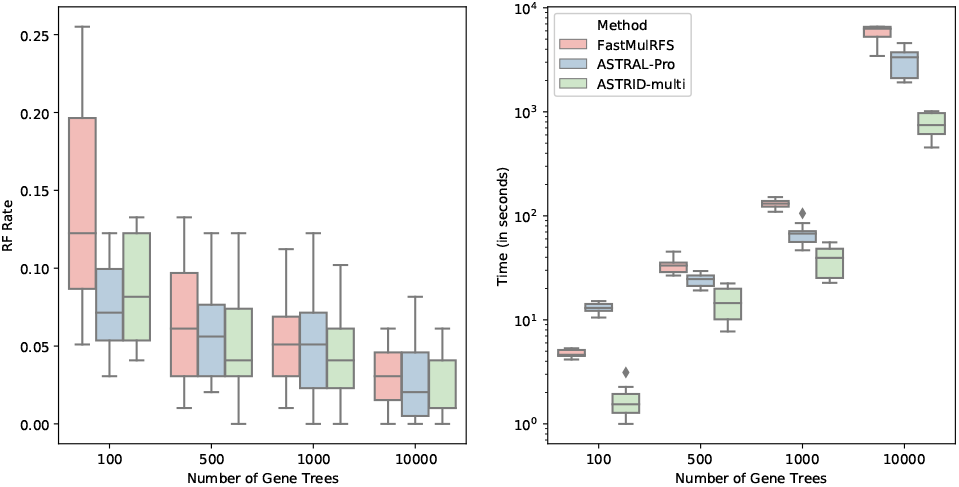
*Experiment 1*. Impact of number of genes on tree error (RF error rates) and wall clock running time (seconds); results shown are averaged across 10 replicates. All the data sets have 100 species, gene trees estimated from 100bp alignments (43.1%,43.3%,43.3%,41.2% MGTE), AD=20%, a duplication rate of 5.0 *×* 10^*−*10^ and an equal loss rate. ASTRAL-Pro ran out of memory on some of the 10,000 gene tree replicates; results shown here are for the replicates on which all methods completed.

### Experiment 2

Next we investigated the effects of missing data (Figure 6). Results shown with missing data are very similar to results shown without missing data (i.e., all gene trees have all the species), but ASTRID-multi’s accuracy degrades slightly compared to ASTRAL-Pro. The relative performance with respect to running time also shows the same trends, with ASTRID-multi fastest and FastMulRFS slowest.

**Fig. 6.**
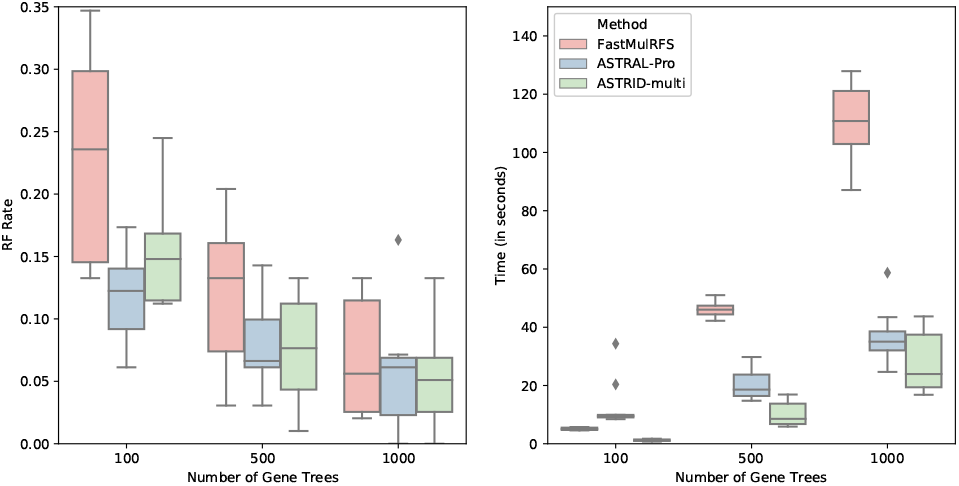
*Experiment 2*. Impact of number of gene trees and missing data on tree error (RF error rates) and wall clock running time (seconds); results shown are for datasets containing 100 species and gene trees estimated from 100bp alignments (43.1%, 43.3%, 43.2% MGTE). All the data sets have AD=20%, a duplication rate of 5.0 *×* 10^*−*10^, and an equal loss rate.

### Experiment 3

We compared methods when analyzing 1000-species 1000-gene datasets, using the Tallis cluster, which allows unlimited time and 256Gb memory (Figure 7). Differences on the five replicates that all methods completed on are very small (and may not be statistically significant), but ASTRAL-Pro showed slightly better accuracy than ASTRID-multi, which was slightly more accurate than FastMulRFS. ASTRID-multi had the advantage when it came to running time, though ASTRAL-Pro was not too much worse (neither exceeded 3 hours for the majority of replicates), and FastMulRFS was by far the slowest with a median time to complete of around 8 hours.

**Fig. 7.**
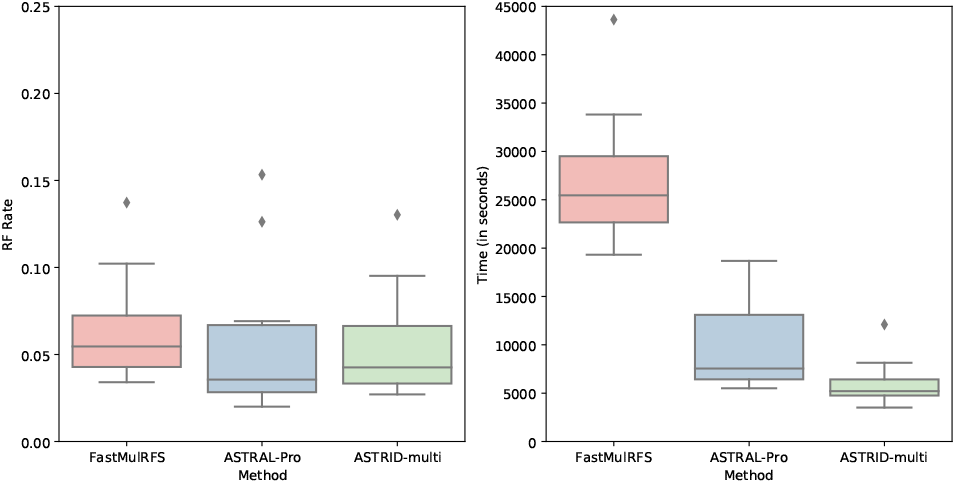
*Experiment 3*. Tree error (RF error rates) and wall clock running time (seconds) on the data set containing 1000 species and 1000 gene trees estimated from 100bp alignments (44.4% MGTE). All the data sets have AD=20%, a duplication rate of 5.0 *×* 10^*−*10^, and an equal loss rate. All methods were run on Tallis.

**Fig. 8.**
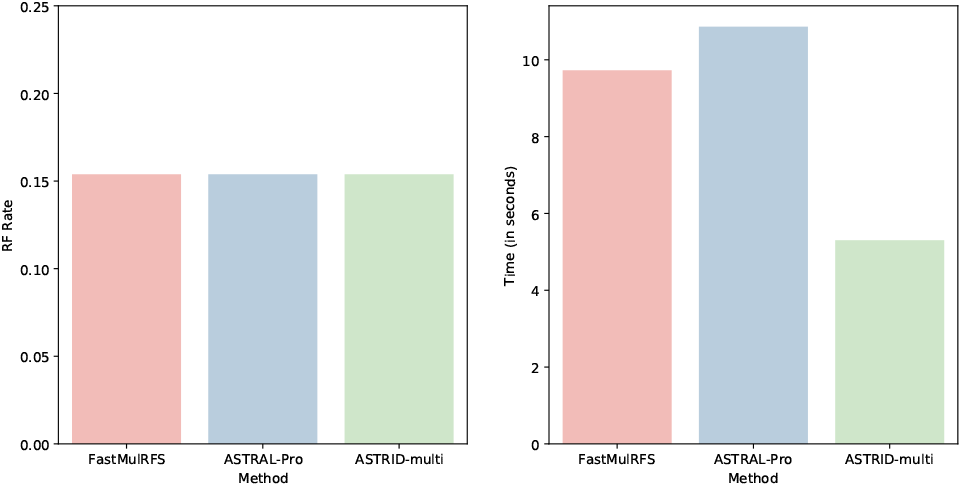
Tree error (RF error rates) and wall clock running time (seconds) on 16-taxon fungal data set with 5351 genes trees from Rasmussen and Kellis [16].

### Failures to Complete

During the course of the study, there were several times that ASTRID-multi or ASTRAL-Pro failed to complete. All these occurred on the Campus Cluster, which has a four hour time limit (Table 2). FastMulRFS always completed, never crashing and also never managing to exceed the four hour time limit. ASTRID-multi crashed twice, both due to memory constraints (once on a replicate with 10,000 gene trees and once on a replicate with a high duplication rate); it also timed out on two high duplication rate replicates. ASTRAL-Pro had the most issues; it timed out on five different replicates (one high duplication rate replicate and four 10,000 gene tree replicates) and crashed on five more due to lack of memory (again, one high duplication rate replicate and four 10,000 gene tree replicates). Notably, ASTRAL-Pro also had difficulty calculating branch lengths with a high number of species, failing to do so on 4 of the 10 replicates with 1000 species. This forced us to re-run these replicates using an argument which suppressed branch length and branch support estimation.

**Table 2.** Failures on the Campus Cluster (4 hr time limit). “Total Failures” includes the total of all failures including out of memory errors (OOM) and timeouts. The campus cluster is heterogeneous; however, at a minimum it has 64 GB of memory. The numbers after GDL is the duplication and loss rate ratio respectively and the number after 10,000 Gene Trees is the loss rate ratio. FastMulRFS is not included as it did not fail to complete in these analyses.

| Method | Experiment | Total Failures | OOM | Timeouts |
| --- | --- | --- | --- | --- |
| ASTRAL-Pro | GDL ($1.0 \times 10^{-9}$ / 0) | 20% | 10% | 10% |
|  | 10,000 Gene Trees (0) | 10% | 10% | 0% |
|  | 10,000 Gene Trees (0.5) | 40% | 0% | 40% |
|  | 10,000 Gene Trees (1) | 30% | 30% | 0% |
| ASTRID-multi | GDL ($1.0 \times 10^{-9}$ / 0) | 30% | 10% | 20% |
|  | 10,000 Gene Trees (0) | 20% | 10% | 0% |

## 4 Discussion and Conclusion

The three methods we compared, ASTRID-multi, ASTRAL-Pro, and FastMulRFS, all had very similar accuracy on these model conditions, but even so some differences can be discerned. Most importantly, when there were substantial differences in accuracy, FastMuLRFS had higher error rates than the other methods; specifically, this occurred on the conditions with high ILS results, high duplication rates, or few genes. In these experiments, there were no model conditions where FastMulRFS was more accurate than the other methods. Thus, in general, although FastMulRFS was in many cases of comparable accuracy to ASTRID-multi and ASTRAL-Pro, typically it is not as reliable as the other methods. The comparison in accuracy between ASTRID-multi and ASTRAL-Pro is therefore of greater interest. In nearly every model condition, the two methods had very similar accuracy, with neither reliably having an advantage over the other. However, small numbers of genes and missing data tended to slightly favor ASTRAL-Pro. Overall, therefore, with respect to accuracy, there is a small advantage to ASTRAL-Pro compared to ASTRID-multi under some conditions, but the difference between the two methods is very small. In general, however, the notable differences in accuracy between methods we observed in these experiments occur only at the extremes (e.g., very high duplication rates, very high ILS, or very few genes). Yet, whether differences in those extreme conditions are important is a matter for discussion. Thus, a basic question that should be addressed is what model conditions are realistic, so that relative performance should be examined for those conditions.

Evaluating methods with respect to use of computational resources shows different trends. Under most conditions, ASTRID-multi is much faster than both ASTRAL-Pro and FastMulRFS, especially for the more challenging datasets where the overall gene tree heterogeneity (which is impacted by MGTE, ILS level, and number of genes) is high. The explanation for this trend is that both FastMuLRFS and ASTRAL-Pro construct a constraint set of allowed bipartitions, which includes at a minimum all the bipartitions from the input gene trees, and the running times for their dynamic programming algorithms (to find optimal solutions within the constrained search space) runs nearly quadratically in the size of the constraint space. In contrast, ASTRID-multi computes a distance matrix, and then computes a tree on the distance matrix using FastME [7]; this calculation is fast except when the duplication rate is very high, and is not impacted by MGTE or ILS (and only minimally impacted by the number of genes). Hence, in nearly all cases, except where there is a very high duplication rate, ASTRID-multi is the fastest of these methods. The methods also differed with respect to failures, caused by memory usage or exceeding the allowed running time: many failures occurred for ASTRAL-Pro, fewer for ASTRID-multi, but none for FastMulRFS. Thus, although ASTRAL-Pro has a small advantage over ASTRID-multi and a somewhat larger advantage over FastMulRFS with respect to accuracy, both are more reliable in terms of failures.

Given the close results in terms of accuracy under most conditions, the choice of method to use may depend on dataset size and computational resources. When memory usage and running time are not of concern, all three methods could be used, and then the bipartitions that appear in all the trees can be considered reliable. When computational resources are more limited, then both ASTRID-multi and ASTRAL-Pro can be used. Finally, on large datasets, if running time is an issue, then ASTRID-multi may be a sufficient method, as it is the fastest and generally close to the accuracy (if not equal to) of ASTRAL-Pro.

